# Prevalence and heterogeneity of antibiotic-resistant genes in *Orientia tsutsugamushi* and other rickettsial genomes

**DOI:** 10.1101/2022.08.17.504356

**Authors:** R. Shyama Prasad Rao, Sudeep D. Ghate, Rajesh P. Shastry, Krishna Kurthkoti, Prashanth Suravajhala, Prakash Patil, Praveenkumar Shetty

## Abstract

Despite a million infections every year and an estimated one billion people at risk, scrub typhus is regarded as a neglected tropical disease. The causative bacterium *Orientia tsutsugamushi*, a member of rickettsiae, seems to be intrinsically resistant to several classes of antibiotics. The emergence of antibiotic-resistant scrub typhus is likely to become a global public health concern. Yet, it is unknown as to how common antibiotic-resistant genes are in *O. tsutsugamushi*, and how variable these loci are among the genomes of rickettsiae. By using the comprehensive antibiotic resistance database, we explored 79 complete genomes from 24 species of rickettsiae for putative antibiotic-resistant loci. There were 244 unique antibiotic-resistant genes in rickettsiae. Both the total and unique antibiotic-resistant genes in *O. tsutsugamushi* were significantly less compared to other members of rickettsiae. However, antibiotic-resistant genes in *O. tsutsugamushi* genomes were more unique and highly variable. Many genes such as resistant versions of *evgS*, and *vanS A/G* were present in numerous copies. These results will have important implications in the context of antibiotic-resistant scrub typhus.

## Introduction

The rapid emergence of antibiotic-resistant bacteria is becoming a global public health crisis (Murray et al., 2022; Ventola, 2015). In 2019, there were an estimated 4.95 million deaths associated with antibiotic resistance worldwide with *Escherichia coli, Staphylococcus aureus, Klebsiella pneumoniae, Streptococcus pneumoniae, Acinetobacter baumannii*, and *Pseudomonas aeruginosa* as the leading/top-six resistant pathogens (Murray et al., 2022).

Scrub typhus – a neglected tropical vector borne zoonotic infectious disease which although is prevalent in “Tsutsugamushi Triangle” of south-east Asian countries, is also increasingly being reported from African and south American countries (Bonell et al., 2017; Chakraborty and Sarma, 2017; Jiang and Richards, 2018; Walker, 2016; Xu et al., 2017). Further, the occurrences of scrub typhus cases are also increasing. For example, in China, the overall incidence has increased sharply from 0.09/100,000 population in 2006 to 1.6/100,000 population in 2016 (Li et al., 2020). In particular, there was a 20-fold increase in infections (n = 27,838) in Yunnan Province between 2006-2017 (Peng et al., 2022). A four-fold increase in infection was seen in South Korea between 2001-2013 with disproportionately more infections in women and older people, and infections mostly occurring during October and November (Lee et al., 2015). Mortality varies widely with a median of 1.4% for treated and 6% for untreated scrub typhus (Bonell et al., 2017; Taylor et al., 2015). Nonetheless, scrub typhus has a high disease burden. It threatens an estimated one billion people globally, and causes illness in one million people each year (Chakraborty and Sarma, 2017; Jiang and Richards, 2018; Xu et al., 2017). The urbanization of scrub typhus has also been described (Li et al., 2020; Park et al., 2015). In south-east Asia, scrub typhus is the leading cause of febrile disease after malaria (Yang et al., 2020).

Scrub typhus is caused by *Orientia tsutsugamushi* (formerly *Rickettsia*) – a gram-negative, obligate intracellular bacillus in the family Rickettsiaceae, and is transmitted to humans by larval form (called chiggers) of arthropod vectors (such as *Leptotrombidium akamushi* and *L. deliense*) in the mite family Trombiculidae. While *O. tsutsugamushi* is the most common re-emerging rickettsial infection in India and many other Southeast Asian countries (Chakraborty and Sarma, 2017; Tilak and Kunte, 2019), members of rickettsiae also cause illnesses such as epidemic typhus by *Rickettsia prowazekii*, murine typhus by *R. typhi*, and spotted fevers by other *Rickettsia* spp. (Rolain et al., 1998) Thus, members of rickettsiae are a persistent threat to public health, and therefore command surveillance (Biggs et al., 2016).

Even though nearly a million infections occur every year, scrub typhus is regarded as a neglected tropical disease (Trent et al., 2019), and WHO has labelled it as one of the most underdiagnosed/underreported diseases (Luce-Fedrow et al., 2018). The symptoms include fever with chills, headache, backache, myalgia, rashes, profuse sweating, vomiting, and enlarged lymph nodes (Lu et al., 2021), and in the absence of early and effective treatment, scrub typhus might lead to interstitial pneumonia, acute respiratory distress syndrome, meningoencephalitis, acute kidney injury, disseminated intravascular coagulation, and death (Walker, 2016). With no vaccine available, antibiotics such as chloramphenicol, doxycycline, macrolides (such as azithromycin), quinolones, rifampicin, and tetracyclines are used to treat scrub typhus (Kelly et al., 2017; Lu et al., 2021; Sayed et al., 2018; Trent et al., 2019). While, treatment with antibiotics is effective for most patients (Kelly et al., 2017; Paris and Wangrangsimakul, 2022; Wangrangsimakul et al., 2020), they might have no significant advantage or disadvantage to others with regard to efficacy or safety (Parola et al., 2017; Yang et al., 2020).

However, *O. tsutsugamushi* has been shown to be intrinsically resistant to several classes of antibiotics including the cephalosporins, gentamicin, penicillins, and possibly the fluoroquinolones (Kelly et al., 2017; Tantibhedhyangkul et al., 2010). Further, resistance to doxycycline and tetracycline has also been suggested (Kim et al., 2008; Lu et al., 2021). Thus, antibiotic resistance in *O. tsutsugamushi* is of great concern, and therefore many studies have been exploring the same (Phuklia et al., 2019; Sayed et al., 2018 and the references therein), including the scrub typhus antibiotic resistance trial – START (Paris and Wangrangsimakul, 2022).

The availability of whole genome sequences is greatly enabling the exploration of antibiotic resistance. Based on the search of loci that might contribute to antibiotic resistance, at least 18 such loci have been shown to occur in the genome of *O. tsutsugamushi*. One gene – *gyrA*, for example, was present as a quinolone-resistant form in the genome of all isolates of *O. tsutsugamushi*. Further, it was also shown that at least 13 other genes that were present in the genus *Rickettsia* did not occur in *O. tsutsugamushi* (Kelly et al., 2017). While these are useful revelations, there remain many open questions. For example, (1) How common antibiotic-resistant genes are in *O. tsutsugamushi* and members of rickettsiae? (2) How variable are these loci among the genomes of a species?

Based on 79 complete genomes from 24 species of rickettsiae and by using the comprehensive antibiotic resistance database of antibiotic-resistant loci, we show the patterns of antibiotic-resistant loci in rickettsiae and reveal their great heterogeneity in the genomes of *O. tsutsugamushi*.

## Materials and Methods

### Acquisition of sequences

The genome sequences were downloaded from the NCBI website (https://www.ncbi.nlm.nih.gov/, last accessed on 07-05-2022). Only the complete genome sequences were used. There were 79 complete genomes from 24 species of rickettsiae, including eight sequences from *O. tsutsugamushi* (Table 1).

**Table 1.** Antibiotic-resistant genes in *O. tsutsugamushi* and other rickettsial genomes.

| # | Species | # of genomes | # of unique ARO<sup>^</sup> | # of ARO<sup>&</sup> | # of unique ARO<sup>+</sup> |
| --- | --- | --- | --- | --- | --- |
| 1 | *Ca. P. trachydisci* | 1 | 65 | 90 | 14 |
| 2 | *O. tsutsugamushi* | 8 | 72 (47-53) | 72.9 (65-86) | 20 |
| 3 | *R. africae* | 1 | 77 | 102 | 0 |
| 4 | *R. akari* | 1 | 68 | 88 | 6 |
| 5 | *R. amblyommatis* | 3 | 83 (79-80) | 107 (107-107) | 3 |
| 6 | *R. asiatica* | 1 | 80 | 107 | 6 |
| 7 | *R. australis* | 1 | 74 | 98 | 5 |
| 8 | *R. bellii* | 3 | 82 (69-74) | 100 (99-101) | 5 |
| 9 | *R. canadensis* | 2 | 60 (59-59) | 78 (77-79) | 3 |
| 10 | *R. conorii* | 6 | 93 (75-78) | 99 (99-99) | 0 |
| 11 | *R. japonica* | 14 | 81 (79-80) | 104.1 (104-105) | 0 |
| 12 | *R. massiliae* | 1 | 77 | 101 | 1 |
| 13 | *R. monacensis* | 1 | 64 | 87 | 3 |
| 14 | *R. montanensis* | 1 | 76 | 102 | 2 |
| 15 | *R. parkeri* | 2 | 80 (74-75) | 102 (101-103) | 0 |
| 16 | *R. peacockii* | 1 | 74 | 96 | 1 |
| 17 | *R. philipii* | 1 | 75 | 99 | 2 |
| 18 | *R. prowazekii* | 10 | 67 (58-62) | 75.3 (74-79) | 4 |
| 19 | *R. rhipicephali* | 2 | 78 (71-72) | 96.5 (96-97) | 4 |
| 20 | *R. rickettsii* | 11 | 80 (72-73) | 96.5 (96-98) | 2 |
| 21 | *R. slovaca* | 2 | 77 (76-76) | 101 (100-102) | 0 |
| 22 | *R. sp. MEAM1* | 1 | 62 | 85 | 1 |
| 23 | *R. tillamookensis* | 1 | 78 | 112 | 6 |
| 24 | *R. typhi* | 4 | 59 (59-59) | 75 (75-75) | 4 |
|  | All | 79 | 244 | 7291 | 92 |

<sup>^</sup>Within species (min-max per genome), <sup>&</sup>Average per genome (min-max), <sup>+</sup>Among all, ARO – Antibiotic resistance ontology.

### Identification of antibiotic-resistant genes

The comprehensive antibiotic resistance database (CARD) was used for the identification of antibiotic-resistant genes. While numerous databases for resistance determinants exist (Doster et al., 2019; Feldgarden et al., 2021; Hendriksen et al., 2019), CARD is perhaps the most comprehensive one. The CARD is a curated resource of over 4336 antibiotic resistance ontology (ARO) terms covering resistance mechanisms from over 2923 known antimicrobial resistance (AMR) determinants/genes and additional 1304 resistance variant mutations (Alcock et al., 2020). The CARD web interface (https://card.mcmaster.ca/analyze/rgi) can quickly identify putative antibiotic-resistant genes based on numerous approaches such as BLAST, sequence alignment, regular expressions (RegEx), hidden Markov models (HMMs), and/or position-specific SNPs (Hendriksen et al., 2019). Each rickettsiae genome sequence was submitted to CARD’s resistance gene identifier (RGI) tool to generate annotation based on perfect, strict, and loose paradigm, and complete gene match criteria for the identification of putative antibiotic-resistant genes (Her et al., 2021; Kent et al., 2020; Zhang et al., 2022).

### Data/statistical analyses

The average and unique number of antibiotic-resistant genes were enumerated for each species. The extent of overlap of genes among different species/groups were represented using a Venn diagram, and visualized using heat map and clustering. The ggvenn() and heatmap.2() functions were used in R, and Bray–Curtis dissimilarity and Ward’s method were used for clustering. A Fisher’s test was used, for example, to check whether the difference in the proportions of genes in two species/groups was significantly different (Agresti, 2018). An unpaired t-test (two-tailed, unequal variance) was used where relevant. The extent of overlap, for example, of genes between two sets, was quantified using overlap coefficient (Vijaymeena and Kavitha, 2016). The data handling/analyses were done in Python and Microsoft Excel.

## Results

### Antibiotic-resistant genes in O. tsutsugamushi and rickettsiae

There were a total of 7291 putative antibiotic-resistant genes in 79 complete genomes of rickettsiae (Table 1 and S1). Altogether, there were 244 unique antibiotic-resistant genes (Table S2). The average number of antibiotic-resistant genes per species ranged from a minimum of 72.9 (SD ±7.7, range 65-86) in *O. tsutsugamushi* to a maximum of 112 in *R. tillamookensis* (Table 1). Compared to rickettsiae at 94.5 (±7.7, 74-112), the average number of antibiotic-resistant genes in *O. tsutsugamushi* was significantly less (p = 2.1E-05, t-test). The number of unique antibiotic-resistant genes within a species ranged from a minimum of 59 in *R. typhi* to a maximum of 93 in *R. conorii*. However, the sets of unique antibiotic-resistant genes were variable among the genomes of a species. In *R. conorii*, for instance, they ranged from 75 to 78 in any one genome, indicating a clear non-overlap of a number of genes. In fact, compared to rickettsiae at 71.6 (±7.7, 58-80), the average number of unique antibiotic-resistant genes per genome in *O. tsutsugamushi* at 49.0 (±2.1, 47-53) was significantly less (p = 6.6E-21). However, as a species, *O. tsutsugamushi* had 72 unique antibiotic-resistant genes.

### Comparison of antibiotic-resistant genes among rickettsiae

A heat map and clustering of 244 antibiotic-resistant genes present among rickettsiae showed that only 23 (9.4%) were common to all 24 species (Fig. 1A and B). The three species namely *O. tsutsugamushi*, *Ca. Phycorickettsia trachydisci*, and *R. belli* formed a close cluster, the typhus fever causing species *R. typhi* and *R. prowazekii*, and a few others formed another sub-cluster, while the spotted fever causing species *R. rickettsii* and *R. conorii*, and the rest formed a lager outer cluster. It may be noted that some 20 antibiotic-resistant genes in *O. tsutsugamushi* (Fig. 1A, and Table 1 and S2) were unique as they were not present in any other rickettsial species. The *Ca. Phycorickettsia trachydisci* had the next highest number of 14 unique antibiotic-resistant genes. Amongst *O. tsutsugamushi* and *R. rickettsii*, *R. typhi*, and *R. prowazekii*, there were 28 common genes (average overlap coefficient of 41.4%), whereas a large set of 17 genes such as *adeS* and *C. difficile gyrA* seemed to be specific to *Rickettsia* (Fig. 1C and Table S2). It may be noted that the version of *gyrA* gene that confers resistance to fluoroquinolones was present in *O. tsutsugamushi* as *A. baumannii gyrA*, whereas all other species had *C. difficile gyrA* (Table S2). The percentages of antibiotic-resistant genes based on the mechanism of resistance are shown in Fig. 1D. While the proportion is higher in efflux category and lower in antibiotic inactivation category for *O. tsutsugamushi* compared to rickettsiae, the difference was not statistically significant (p = 0.29, Fisher’s test).

**Fig. 1.**
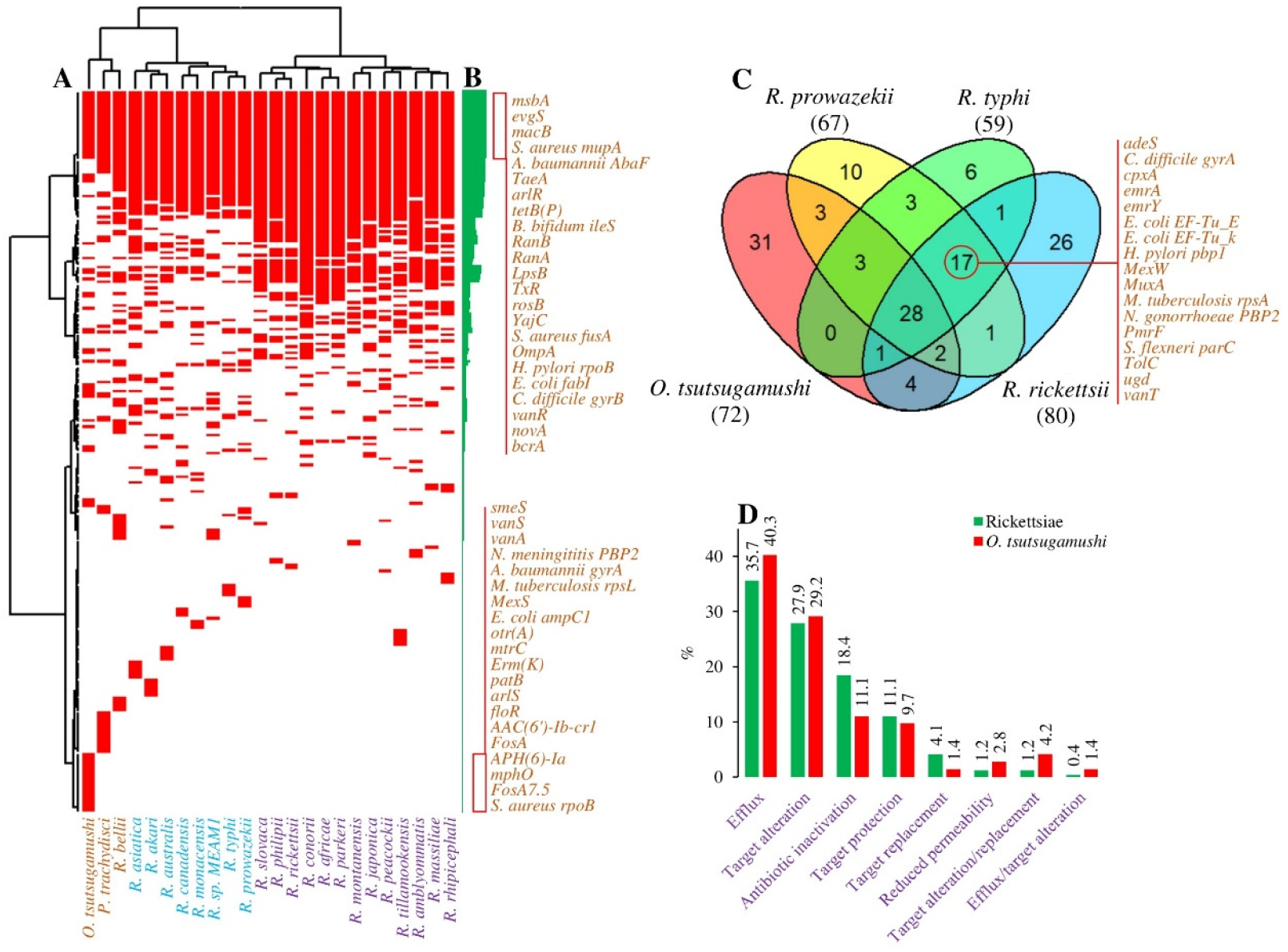
Antibiotic-resistant genes among rickettsiae. (A) Of the 244 potential antibiotic-resistant genes, just 23 (9.4%) are present in all rickettsial species. *O. tsutsugamushi* has a large number of unique antibiotic-resistant genes. Panel (B) shows frequency (max 24 species) for each gene. (C) Overlap of antibiotic-resistant genes among *O. tsutsugamushi* and three key *Rickettsia* species. (D) Percentage of antibiotic-resistant genes based on mechanism of resistance. See Table S1 for the complete list of antibiotic-resistant genes.

### Heterogeneity of antibiotic-resistant genes in O. tsutsugamushi

Where multiple (four or more) genome sequences available, we looked at the variability of antibiotic-resistant genes among the genomes in five species – *O. tsutsugamushi, R. japonica, R. prowazekii*, *R. rickettsii*, and *R. typhi* (*R. conorii* was ignored due to some plausible annotation issue in one of the sequences). Of the 147 antibiotic-resistant genes amongst these five species, 62 (42.2%) genes were variable in the genomes of any one of the species (Fig. 2A). For instance, the *E. coli ampH* gene, while present/absent in all genomes of other species, was present only in one out of eight genome sequences of *O. tsutsugamushi*. In fact, of the 72 antibiotic-resistant genes in *O. tsutsugamushi*, 38 (52.8%) were variable, and that percentage was significantly higher (p = 4.9E-05) than the next highest of 19.4% in *R. prowazekii* (Fig. 2B). There was no variability of antibiotic-resistant genes in *R. typhi* genome sequences.

**Fig. 2.**
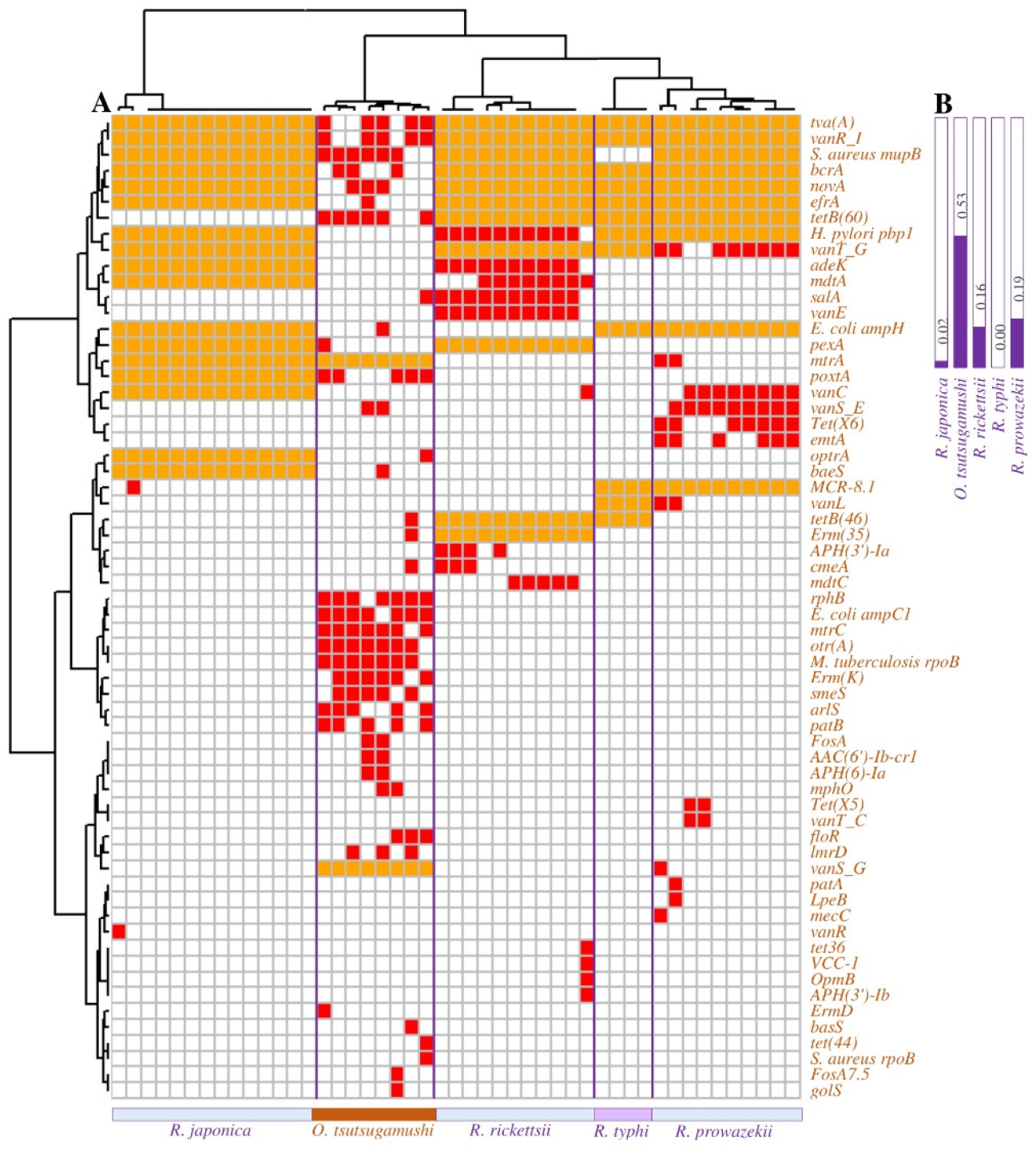
(A) Heterogeneity of antibiotic-resistant genes among genomes of five species of rickettsiae. The antibiotic-resistant genes that are not present in all genomes within a species are shown in red. (B) A large proportion (≈0.53) of *O. tsutsugamushi* antibiotic-resistant genes are heterogeneous.

Further, of the 244 antibiotic-resistant genes amongst rickettsiae, 53 (21.7%) genes were present in multiple (two or more) copies in any one of the species/genomes (Fig. 3A). Of these, 32, 14, and three genes were present up to a maximum of two, three, or four copies, respectively; whereas remaining four genes namely *A. baumannii AbaF*, *evgS*, *vanS A*, and *vanS G* were present in 10 or more copies in some genomes (Fig. 3B and C). For example, there were two copies of *A. baumannii AbaF* in *O. tsutsugamushi*, but 16 copies in *R. tillamookensis*. Further, *O. tsutsugamushi* had, on average, three copies of *vanS A*; however, it varied from one to 10 copies in individual genome sequences (Fig. 3C).

**Fig. 3.**
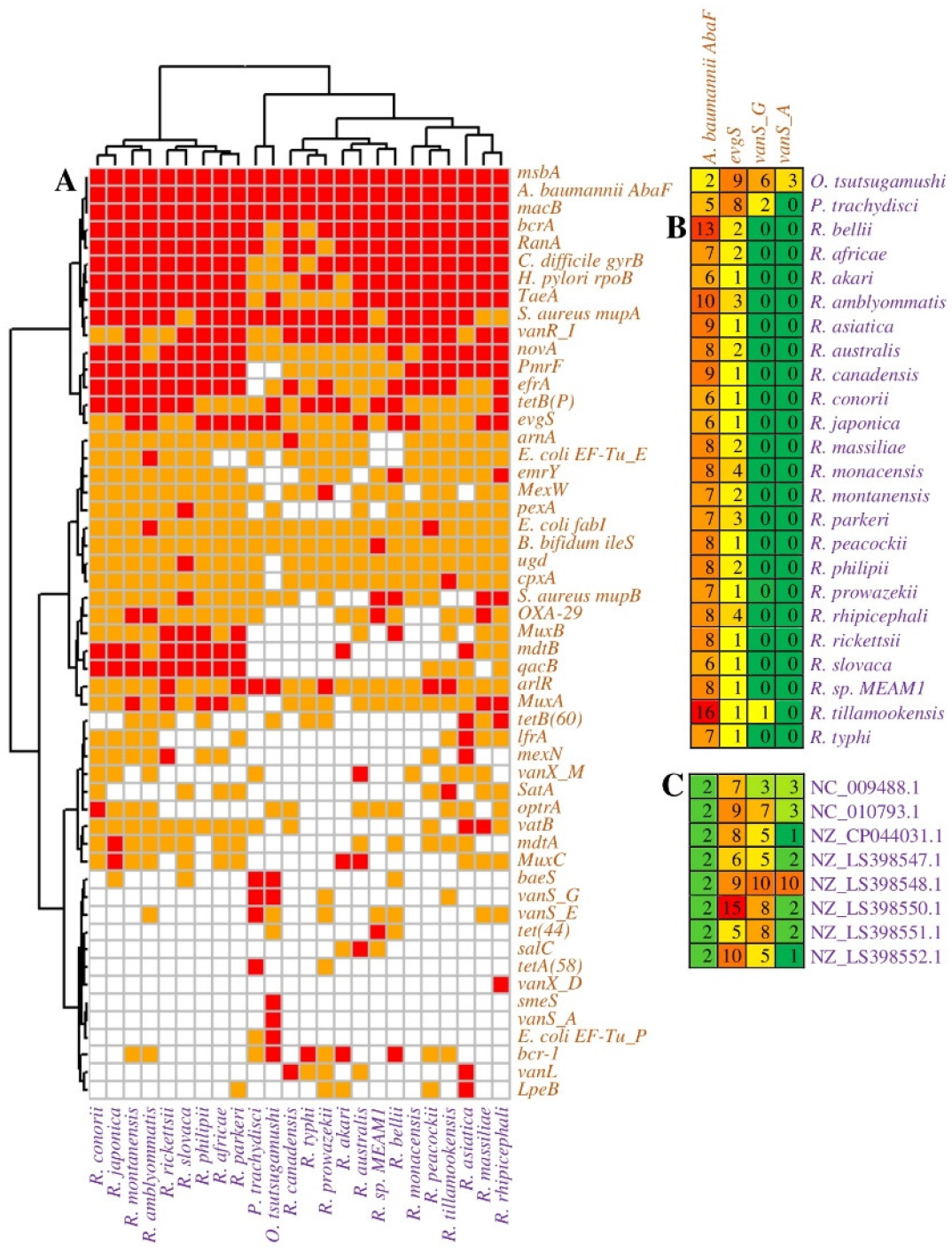
Heterogeneity of antibiotic-resistant genes among species of rickettsiae. (A) The antibiotic-resistant genes that are present in multiple (two or more) copies within a species are shown in red. The number of copies for top four antibiotic-resistant genes (which are present in 10 or more copies in any one genome) are shown for (B) different species of rickettsiae and (C) different genomes of *O. tsutsugamushi*.

## Discussion

The emergence of resistance to antibiotics is the most challenging issue in the treatment of bacterial infections (Uddin et al., 2021). Antibiotic-resistant infections are widespread across the globe (Ventola, 2015; Zhang et al., 2022). Most bacteria might contain some form of antibiotic-resistant genes such as resistance plasmids or efflux pumps that might remain functionally silent until sufficiently challenged with selection pressure (Nikaido, 2009). Given the growing number of cases, scrub typhus is emerging as a global public health threat (Devasagayam et al., 2021). As the scrub typhus is intrinsically resistant to many antibiotics (Tantibhedhyangkul et al., 2010), it might pose even a greater danger.

In this work, we showed that all rickettsial species contain numerous putative antibiotic-resistant loci. For instance, there were numerous variants of *rpoB* which confers resistance to rifampicin. Similarly, there were many other putative loci such as *ampC1*, *ampH*, and *PBP2* which confer resistance to beta-lactam, and *pbp1/2/3* which confer resistance to amoxicillin. Numerous rickettsial species were experimentally shown to be resistant to rifampicin. Likewise, it was also known that beta-lactams and aminoglycosides are not effective, and amoxicillin, gentamicin, and co-trimoxazole have poor sensitivity in treating rickettsial diseases (Rolain et al., 1998).

Although the number of antibiotic-resistant loci were significantly less compared to other rickettsiae, they were more unique and highly variable in *O. tsutsugamushi* genomes. In comparison, there was no inter-genome variability of antibiotic-resistant loci in *R. typhi*. The *O. tsutsugamushi* is known to have one of the most highly repeated bacterial genomes sequenced (Cho et al., 2007). As against 2,179 potential protein-coding loci, the number of putative antibiotic-resistant loci is very small in *O. tsutsugamushi*. However, given that the genome contains CRISPR-like elements including more than 400 transposases, 60 phage integrases, and 70 reverse transcriptases (Cho et al., 2007), the *O. tsutsugamushi* has enough gears to tinker its genome under selection. The *O. tsutsugamushi* was known to have high antigenic diversity. In India, for instance, Kato-like (NZ_LS398550.1) strains predominate (61.5%), followed by Karp-like (NZ_LS398548.1) strains (27.7%), and Gilliam and Ikeda strains (Varghese et al., 2015). Further, *O. tsutsugamushi* was also known to undergo genetic recombination among diverse genotypes (Kelly et al., 2017; Kim et al., 2017). The extent of diversity and heterogeneity of putative antibiotic-resistant loci in *O. tsutsugamushi* gives a hint that there is potential to gain antibiotic resistance under selection.

Microbes are said to harbour a ‘silent reservoir’ of antibiotic-resistant genes that is thought to contribute to the emergence of multidrug-resistant “superbugs” through horizontal gene transfer (Kent et al., 2020). While horizontally acquired antibiotic-resistant genes via plasmids are common (Bennett, 2008; van Hoek et al., 2011), this may not be common in rickettsiae as it is said that intracellular lifestyle restricts the opportunity for lateral gene transfer (Vanrompay et al., 2017). However, it should be noted that *O. tsutsugamushi* genome has 359 tra genes for components of conjugative type IV secretion systems which play important role in horizontal gene transfer, and other rickettsiae too, such as *Rickettsia felis*, have numerous plasmid-encoded tra genes (Cho et al., 2007).

It is important to note that many genes such as resistant versions of *AbaF*, *evgS*, and *vanS A/G* are present in multiple copies in rickettsiae. The *AbaF*, for instance, is a well-known efflux pump that is involved in antibiotic resistance (Abdi et al., 2020). Any perturbations under antibiotics such as mutations leading to increased expression of efflux pumps may impart antibiotic resistance (Nikaido, 2009; Salini et al., 2022).

To mention the limitations of this study, we used only the complete genomes, and not more numerous partial/incomplete genome sequences. More importantly, the antibiotic-resistant loci were loose hits in the CARD database. Finally, as this is a bioinformatic analysis, like others (Her et al., 2021), we do not make any experimental validations.

In conclusion, we showed that there is a wide diversity of putative antibiotic-resistant genes in rickettsiae. Further, they were more unique and highly variable in *O. tsutsugamushi* genomes. Given sufficient selection pressure/challenge, *O. tsutsugamushi* and other rickettsiae have plenty of potential loci, such as resistant versions of *gyrA* and efflux pump *AbaF*, to develop antibiotic resistance. Thus, surveillance of antibiotic resistance should be a priority to avoid a global public health crisis.

## Funding and Acknowledgments

This work did not receive any specific funding.

## Statement of Ethics

The work is in compliance with ethical standards. No ethical clearance was necessary.

## Data Availability

The sequence data used in this work were obtained from NCBI. The relevant derived data are given in supplemental Tables S1 and S2.

## Author Contributions

RSPR, SDG, and RPS planned the work, and RSPR and SDG performed the work and wrote the manuscript. All authors contributed intellectually, and reviewed the manuscript.

## Supplemental Information

Supplemental information for this article is available online.

